# Noradrenaline modulates decision urgency during sequential information gathering

**DOI:** 10.1101/252932

**Authors:** Tobias U. Hauser, Michael Moutoussis, Nina Purg, Peter Dayan, Raymond J. Dolan

## Abstract

Arbitrating between timely choice and extended information gathering is critical in effective decision making. Aberrant information gathering behaviour is said to be a feature of psychiatric disorders such as schizophrenia and obsessive-compulsive disorder. We know little about the neurocognitive control mechanisms that drive such information gathering. In a double-blind placebo-controlled drug study with 60 healthy humans (30 female), we examined the effects of noradrenaline and dopamine antagonism on information gathering. We show that modulating noradrenaline function with propranolol leads to decreased information gathering behaviour and this contrasts with no effect following a modulation of dopamine function. Using a Bayesian computational model, we show sampling behaviour is best explained when including an urgency signal that promotes commitment to an early decision. We demonstrate that noradrenaline blockade promotes the expression of this decision-related urgency signal during information gathering. We discuss the findings with respect to psychopathological conditions that are linked to aberrant information gathering.

**Significance Statement:** Knowing when to stop gathering information and commit to an option is non-trivial. This is an important element in arbitrating between information gain and energy conservation. In this double-blind, placebo-controlled drug study, we investigated to role of catecholamines noradrenaline and dopamine on sequential information gathering. We found that blocking noradrenaline led to a decrease in information gathering, with no effect seen following dopamine blockade. Using a Bayesian computational model, we show that this noradrenaline effect is driven by an increased decision urgency, a signal that reflects an escalating subjective cost of sampling. The observation that noradrenaline modulates decision urgency suggests new avenues for treating patients that show information gathering deficits.

## Introduction

A rushed decision without the benefit of sufficient information can be detrimental especially if an outcome has non-trivial consequences (e.g., buying a house). By contrast, excess information gathering for trivial decisions (e.g., which toothpaste to buy) can waste time and energy. This dilemma in arbitrating between extended information gathering versus making a more time-efficient, albeit less well informed, choice is often referred to as a speed-accuracy trade-off (Martin and Müller, 1899; Henmon, 1911).

This tradeoff is well captured in sequential information gathering tasks (Huq et al., 1988; Chamberlain et al., 2007). Several psychiatric disorders including schizophrenia are consistently associated with a reduced propensity to gather information, often termed as ‘jumping-to-conclusions’. Such behaviour is found in patients (Fear and Healy, 1997; Moutoussis et al., 2011; e.g., Dudley et al., 2016), but is also a feature of prodromal states (Rausch et al., 2016) and first-degree relatives (Van Dael et al., 2006). This contrasts with patients diagnosed as having obsessive-compulsive disorder (OCD) who tend to gather more information (Volans, 1976; Fear and Healy, 1997; Pélissier and O’Connor, 2002; Voon et al., 2016; Hauser et al., 2017c, 2017b), although not ubiquitously so (Chamberlain et al., 2007; Jacobsen et al., 2012; Grassi et al., 2015).

The neurocognitive control mechanisms that drive these effects is unknown. Using Bayesian computational modelling, we recently showed that one key contributor is a decision urgency signal that promotes timely decisions (Hauser et al., 2017c, 2017b). The neural processes that modulate such urgency signals in sequential information gathering remain unknown. In this study, we examined the role of catecholaminergic neuromodulators in information gathering. Dopamine is implicated in the genesis of schizophrenia (Laruelle, 2013), but previous studies found inconsistent results about its role in jumping to conclusions (Menon et al., 2008; So et al., 2010, 2012; Ersche et al., 2011; Andreou et al., 2013, 2015; Ermakova et al., 2014). Other evidence points to a more critical role for noradrenaline, including role in signalling uncertainty (Aston-Jones and Cohen, 2005; Yu and Dayan, 2005; Dayan and Yu, 2006), suggesting this neuromodulator might play a role during information gathering.

Here, we assessed the contributions of both dopamine and noradrenaline to information gathering in 60 healthy subjects. Using a double-blind placebo-controlled design, we examined the effects of catecholaminergic antagonists with a high specificity for either dopamine (amisulpride) or noradrenaline (propranolol). We show that blocking noradrenaline modulates information gathering in an information sampling task, whereas dopamine antagonism had no significant behavioural consequence. Using computational modelling, we demonstrate that the effect of propranolol is best accounted for via alteration of an urgency signal.

## Materials and Methods

### Subjects

We conducted a double-blind, placebo-controlled, between-subjects study involving three groups of 20 subjects each. Subjects were randomly allocated to one of the three groups (ensuring an equal gender balance) after excluding those who met the following exclusion criteria: a history of psychiatric or neurological disorder, regular medication (except contraceptives), current health issues, any drug allergies. The groups did not differ in their mood (PANAS) (Watson et al., 1988), intellectual ability (WASI abbreviated version) (Wechsler, 1999) or age (Table 1). Data from the same sample has been reported previously (Hauser et al., 2017a) that addressed a different topic. UCL research ethics approved this study and all subjects provided written informed consent.

**Table 1:** Drug group characteristics. The drug groups were matched for an equal gender balance and did not differ in their age, intellectual abilities (WASI) or their affect (PANAS). mean±SD.

|  | **placebo** | **propranolol** | **amisulpride** |  |
| --- | --- | --- | --- | --- |
| gender (m/f) | 10/10 | 10/10 | 10/10 |  |
| IQ | 112.45±12.22 | 118.75±8.55 | 114.60±11.77 | F(2,57)=1.70 , p=.191 |
| age | 24.50±4.16 | 23.15±4.31 | 22.35±2.21 | F(2,57)=1.74 , p=.185 |
| positive affect | 29.22±10.47 | 27.15±7.75 | 27.80±8.12 | F(2,57)=.286 , p=.752 |
| negative affect | 11.45±2.37 | 11.95±4.87 | 11.25±1.92 | F(2,57)=.236 , p=.790 |

### Drug groups

To assess the effects of neurotransmitters dopamine and noradrenaline on information gathering, we used three different drug conditions. The noradrenaline group received 40mg of propranolol (β-adrenoceptor antagonist). The dopamine group received 400mg of the D2/3 antagonist amisulpride. We selected these drugs because they have an affinity for the targeted neurotransmitter with a high specificity. The dopamine group received the active drug 120 minutes prior to the task and an additional placebo 30 minutes after the first drug. The noradrenaline group first received a placebo, and after 30 minutes the active drug. A third, placebo group received placebo at both time points. This schedule aligned with procedures used in previous studies investigating the effects of dopamine or noradrenaline on cognition (Silver et al., 2004; Gibbs et al., 2007; e.g., De Martino et al., 2008; Hauser et al., 2017a; Kahnt and Tobler, 2017).

### Experimental design: information gathering task

We examined sequential information gathering using a modified version of an information sampling task (Clark et al., 2006; Hauser et al., 2017c, 2017b). In each game, subjects saw 25 covered cards (depicted by gray squares, Fig. 1A) and had to decide whether the majority of cards comprised yellow or blue (colours varied across games). The subjects were allowed to open as many cards as they wished before committing to one of the two colours.

The first 15 games belonged to a ‘fixed’ condition, in which there was no external cost for sampling more information. Subjects received 100 points for correct decisions and lost 100 points for incorrect decisions, independent of the number of cards opened or the time spent on task prior to decision. The second 15 games belonged to a ‘decreasing’ condition, in which information gathering was accompanied by a reduction in potential wins: starting from a potential win of 250 points, opening each card led to a 10-point reduction in wins (e.g. win after 5 opened cards: 250 – 5 * 10 = 200 points). Incorrect decisions were punished with −100 points irrespective of the amount of prior sampling.

Before the first game, subjects performed a single practice game to familiarise themselves with the task. After each decision, subjects were informed about their winnings and then directly moved to the next game. The game sequences were selected so that 10 games in each condition were relatively difficult with a generative probability close to 50% (similar to that in the original information sampling task; (Clark et al., 2006)). An additional 5 sequences were easier with a clearer majority (generative probabilities of a binomial process p~.7) to allow for a broader variability in information gathering (order of sequences was randomised).

### Statistical analysis

In this study we tested whether blockade of dopamine or noradrenaline function impacted on information gathering behaviour. The number of draws before a decision is a good indicator for the amount of information that a subject opts to collect before making a decision. We thus analysed this behavioural metric using a repeated-measures ANOVA with the between-subject factor group (propranolol, amisulpride, placebo) and the within-subject factor condition (fixed, decreasing). Effects were further assessed using independent-sample t-tests. Effects sizes are indicated using partial eta-squared and Cohen’s d. As secondary measures, we also assessed whether a group won more points or was less accurate in their decision making (i.e. how often subjects decided for a current minority of cards) using the same statistical procedures.

### Computational modelling

To understand cognitive mechanisms of how decisions arise, and to probe deeper onto how drugs may affect these cognitive processes, we used a Bayesian computational model that we previously developed and validated for this task (Hauser et al., 2017c, 2017b). In brief, the winning model assumes that at each state of the game subjects arbitrate between three actions: deciding for yellow, deciding for blue, continuing with sampling (non-deciding). This arbitration is based on a decision policy, which in turn is based state-action Q-values (Watkins, 1989) of each option. The Q-values for deciding in favour of either colour are computed as the outcome of a correct/incorrect decision, weighted by their inferred likelihood (i.e. ‘how likely am I to win if I decide for yellow now, given the cards I have opened so far?’). The action-value for continuing sampling indexes the value of future states (using backwards induction), plus a subjective cost per step. Notably the latter captures an urgency to decide that arises as sampling continues.

Here, we reiterate the key equations of the winning model for completeness. Please see Hauser et al. (2017c) for a more detailed description and discussion (also cf Moutoussis et al., 2011; Hauser et al., 2017b).

Our model assumes that subjects try to infer the colour that forms the majority of cards, based on the cards seen so far. This means that the subjects infer the probability that the majority of cards belongs to a particular colour (e.g. yellow (*y*))*P(MY|n_y_,N*), where *MY* depicts a majority of yellow cards, *n_y_* the number of opened yellow cards, and *N* the total number of opened cards.

*P(MY)* is fully determined as soon as 13 or more of the opened cards belong to one colour (of a total number of cards: *N_tot_*=25). This can be inferred by calculating the probability of the majority of cards being yellow, given a specific generative probability *q* (proportion of yellow and blue cards in the machinery that produces the sequence), weighted by the likelihood of this generative probability based on the currently seen cards:

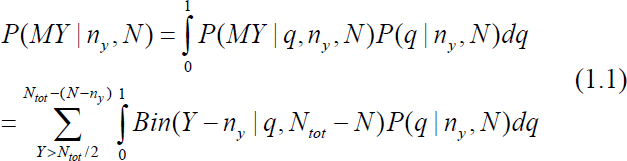

The first expression is a binomial of getting *Y-n_y_* yellow draws out of *N_tot_-N*, given generative probability *q*. The second expression is the probability of the *q* being the generative probability. This can be calculated using Bayes rule:

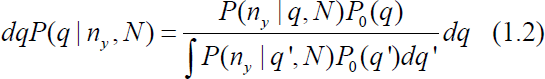

We assume that the probability of *n* yellow draws of *N* total draws follows a binomial distribution and that the prior belief about the generating *q* follows a beta distribution (conjugate prior) with the parameters *α* and *β* (here *α*=1, *β*=1).

The beliefs about the majority of cards are subsequently translated into action-values. The action value of choosing *Y*, (*Q(Y)*), is the product of reward/cost of choosing the right or wrong option (*R_cor_*, *R_inc_*) and the success-probabilities of these actions. *Q(B)* is calculated analogously.

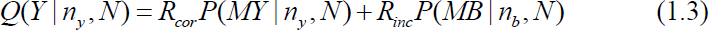

The rewards of correctly (*R_cor_*) and incorrectly declaring (*R_inc_*) can be cast in different ways. According to the objective instructions, in the fixed condition *R_cor_* is set to 100 and *Rinc* to −100. For the decreasing condition, we compared two different formulations. In our main model (‘subjective’), we also kept *R_cor_* constant in the decreasing condition. This was done so that the subjective costs (*c_s_*, cf below) soak up the subjectively perceived overall costs, i.e. a combination of externally imposed and internally generated costs. This way, we can investigate the subjectively perceived total costs. Alternatively, we formulated an ‘objective’ costs model, where *R_cor_* changes as a function of step (250, 240, 230, …), as set up in the task. This ‘objective’ model only differed in the decreasing, but not in the fixed condition. Additionally, *R_inc_* was kept at −100 for all models and conditions (as per instructions). For the simulation of an optimal model (green diamonds in Fig. 1), we used the objective model and assumed no additional costs-per-step.

The action value of not deciding (*Q(ND)*) computes the value of future states in terms of the future action values and their probabilities. Additionally, a cost per step is imposed that assumes that there are internal (and external) costs that emerge when continuing with sampling (urgency signal). *Q(ND)* is calculated using backward induction to solve the Bellman equation, using state values *V(s’)* and a cost per step *c_s_*:

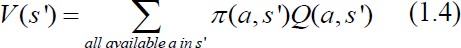

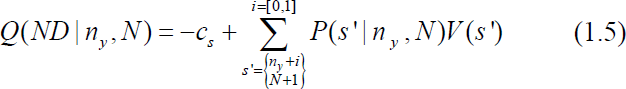

The probability of reaching state *s’* and seeing *i* new yellow items is based on the current belief state, which in turn is mainly determined by the current evidence *n_y_*, *N*. Thus:

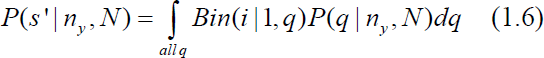

The choice policy *π* for the state-action space is specified as the following softmax function with decision temperature parameter *τ* and irreducible noise (lapse rate) parameter *ξ* (cf Guitart-Masip et al., 2012):

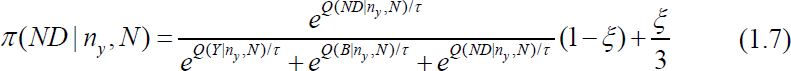

### Decision urgency: nonlinear sigmoidal cost function

We compared two different possible cost functions: fixed cost per sample (‘linear’) vs a model in which the costs per sample increased according to a sigmoid function (‘nonlinear’, cf below). The latter could capture the possibility that subjects felt an increasing urgency (Cisek et al., 2009) to decide, for instance if it becomes increasingly annoying to gather more samples with potentially little informational content and waste time, similar to previous reports that show that costs increase nonlinearly (Drugowitsch et al., 2012; Murphy et al., 2016).

We implemented the nonlinear cost function as a sigmoid, where the cost per step *c_s_* (eq. (1.5)) changes on each step *n* (1, …, 25):

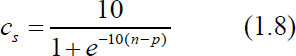

The free impatience parameter *p* describes the indifference point, i.e. at what stage in the game the agent becomes impatient. We set the scaling factor as well as the slope parameter to 10. Model comparison revealed that fixing these parameters led to an equal performance in terms of model fit, and thus outperformed other, more complex models that had these parameters as free parameters. Parameter *p* was independently modelled in the two conditions, allowing for different urgency trajectories.

### Model fitting and model comparison

Using a genetic algorithm implemented in Matlab (Goldberg, 1989), we optimized the parameters to maximise a LogLikelihood for each participant individually. For model comparison, we used summed AIC (Akaike, 1973) and BIC (Schwarz, 1978). The best fitting model was then used for further analyses as reported in the main manuscript.

Similar to our previous findings (Hauser et al., 2017c, 2017b), a subjective costs model (logL: −6239, AIC: 13079, BIC: 13333) clearly outperformed an objective model that incorporates the explicit external costs (logL: −8638, AIC: 17875, BIC: 18129). Further, we found that a model with a nonlinear cost-function outperformed a linear cost model (logL: - 4315, AIC: 9709, BIC: 10133). This means that decision urgency arises in a non-linear, sigmoidal fashion, escalating as sampling continues. This is in line with the sorts of non-linear urgency-signals that are considered to influence a class of perceptual decision making problems (e.g., Churchland et al., 2008; Cisek et al., 2009; Drugowitsch et al., 2012; Murphy et al., 2016).

Further model comparison revealed that a model with only one free parameter *p* (midpoint of a sigmoid) per condition in the costs function achieved the same model fits as more complex versions with additional free parameters, and thus outperformed these models in terms of AIC and BIC (logL: −4365, AIC: 9331, BIC: 9585). Model simulations using this winning model revealed the same behavioural patterns as in the subjects’ actual behaviour (not shown.

To assess how noradrenaline modulated the computational mechanisms underlying information gathering, we focused on a comparison between placebo and propranolol and whether the decision urgency arose at different time points. To do this we used both the model-derived urgency signal as well as the model parameter *p* that modulates its emergence. A group difference for the former was assessed using cluster-extent permutation tests (p<.05, height threshold t=1, 1000 iterations; cf. Hunt et al., 2013; Hauser et al., 2015). This urgency is also critical for modulating dynamic decision thresholds, which indicates how much evidence in favour of one choice option is needed for making a decision (Ratcliff and McKoon, 2008). Consequently, we analysed dynamic decision thresholds which varied over sampling, quantified as the mean evidence difference needed to decide for a simulated agent with subject-specific model parameters.

## Results

### Noradrenaline blockade decreases information gathering

An analysis of the number of draws before declaring a choice, as an index for information gathering, revealed a main effect of group, supporting a significant drug effect on information gathering (Fig. 1B, F(2,57)=4.29, p=.018, η^2^=.131). Follow-up analysis showed this effect was driven by a reduction in information gathering in the noradrenaline compared to a placebo group in both conditions (fixed condition: t(38)=2.55, p=.015, d=.81; decreasing condition: t(38)=2.71, p=.010, d=.86). The dopamine group showed an intermediate effect but did not significantly differ from either group (fixed condition: vs placebo: t(38)=1.53, p=.134, d=.48; vs noradrenaline: t(38)=1.08, p=.286, d=.34; decreasing condition: vs placebo: t(38)=1.47, p=.150, d=.47; vs noradrenaline: t(38)=1.47, p=.149, d=.46). A main effect of condition, but no interaction with group, indicated that subjects across placebo and drug conditions gathered more information in the fixed compared to the decreasing condition (condition main effect: F(1,57)=145.92, p<.001, η^2^=.719; interaction: F(2,57)=.528, p=.592, η^2^=.018).

**Figure 1.**
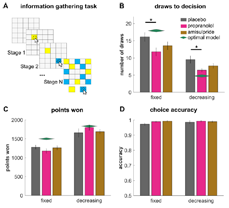
Noradrenaline blockade diminished information gathering. (A) Starting from a fully covered deck of cards, subjects were allowed to open as many cards until they felt ‘certain enough’ to indicate whether they believed the majority of the 25 cards was yellow or blue. In a ‘fixed’ condition, no external costs for sampling applied, but in the ‘decreasing’ condition, a potential win of 250 points reduced by 10 points per uncovered card. (B) Information gathering is decreased consequent on noradrenaline blockade (propranolol), but not consequent on a dopamine perturbation (amisulpride). This increase in impulsivity is consistently observed across both conditions, rendering the statistics of choices in the noradrenaline group closer to those of an optimal model (green diamonds) in the decreasing condition, but further away in the fixed condition. The drug manipulation did not lead to a statistically significant difference in points earned (C), and did not affect choice accuracy (D, the probability of choosing the colour that currently forms the majority of cards at time of decision). * p < .05; mean±1s.e.m.

### No significant drug effects on winnings or accuracy

To test whether the noradrenaline or dopamine drug affected other aspects of performance, we compared the total points won as well as accuracy of decisions. There was no group effect on winnings (Fig. 1C, F(2,57)=.04, p=.961, η^2^=.001) or an interaction (F(2,57)=1.67, p=.198, η^2^=.055), suggesting that the reduced information gathering in noradrenaline was not large enough to impact subjects’ winnings. This absence of a difference in winnings (placebo vs propranolol: fixed condition: t(38)=1.32, p=.195, d=.42; decreasing: t(38)=-1.06, p=.294, d=.33) reflects the fact that most sequences had a generative probability close to 50%. Therefore, a decrease in information/decrease in costs under propranolol were not large enough to percolate through to actual winnings. The groups did not differ in choice accuracy (Fig. 1D, evident in a lack of main effect of group: F(2,57)=1.90, p=.159, η^2^=.063; the absence of an interaction: F(2,57)=.86, p=.428, η^2^=.029). This indicates the drug manipulation did not impact on general motivation to solve the task correctly.

### Propranolol promotes urgency-to-decide

To understand the cognitive processes that drive a lowered information gathering disposition in the noradrenaline group, we fitted a Bayesian computational model (cf. Hauser et al., 2017c, 2017b) to individual subjects’ data. Decision policy as a function of evidence and sampling is depicted in Fig. 2.

**Figure 2.**
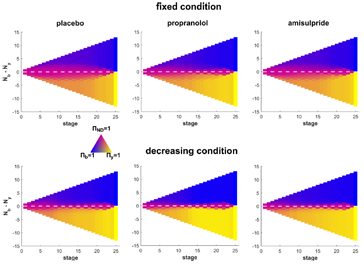
Model policy. The model policy reflects the choice probability for choosing yellow, blue or continuing sampling (pink), depending on the evidence difference (y-axis) and the information gathering stage (x-axis). This depiction shows that subjects are less likely to sample in the decreasing condition in general (less pinkish areas). Also, one can clearly see how a non-decision area is diminished in the propranolol group (middle panels) compared to the placebo group (left panels).

In the model, the speed with which subjects make a decision is modulated by two separate processes: a finite horizon and subjective urgency. The former arises because there is a limit on the number of cards such that whenever 13 cards of the same colour are opened, opening more cards does not provide further information. The latter, however, is not build into the task explicitly (especially in the fixed condition), but is based on a finding that subjects apparently become more liberal in their decision criterion as sampling progresses (Cisek et al., 2009; Drugowitsch et al., 2012; Thura and Cisek, 2014; Murphy et al., 2016). In the model, urgency is captured by a subjective cost for each sample, and we found that it arises in a nonlinear fashion (being low in the beginning but escalates over the course of sampling). The cost arose significantly earlier in the noradrenaline compared to placebo condition in both conditions (Fig. 3, fixed: p=.003, decreasing p=.029) reflecting their lower number of draws.

**Figure 3.**
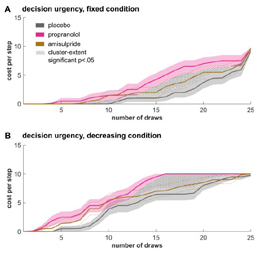
Premature urgency drives noradrenaline-modulated decreases in information gathering. A cost per step signal reflects an emerging urgency to commit to a decision as sampling progresses. This urgency signal arises nonlinearly, escalating significantly earlier in the noradrenaline group in the fixed (A) and the decreasing (B) condition. mean±1s.e.m.

We compared the individually fitted model parameters between groups and found a group effect in the repeated-measures ANOVA (F(2,57)=4.42, p=.016, η^2^=.134) for the parameter *p.* This describes the mid-point of an escalating urgency signal. Subsequent t-tests revealed that the noradrenaline group expressed significantly lower parameter values in both conditions (Table 2; *p_fixed_*: t(38)=2.60, p=.013, d=.82; *p_decreasing_*: t(38)=2.94, p=.006, d=.93), meaning subjective costs/urgency arise significantly earlier in the sampling process. The dopamine group, again, did not differ from other groups (*p_fixed_*: vs placebo: t(38)=-1.30, p=.201, d=.41; vs noradrenaline: t(38)=1.21, p=.232, d=.38; *p_decreasing_*: vs placebo: t(38)=-1.10, p=.279, d=.35; vs noradrenaline: t(38)=1.54, p=.132, d=.48). The choice stochasticity parameter *τ* (F(2,57)=.25, p=.783, η^2^=.009) as well as the lapse rate parameter *ξ* (F(2,59)=1.10, p=.340, η^2^=.037) did not yield significant group differences.

**Table 2:** Model parameter comparison. Group comparison reveals that the propranolol group has significantly lower indifference point parameters for both the fixed (*pf*) and the decreasing (*pd*) condition. This means that their urgency arises earlier in an information gathering process and thus drives them to make more hasty decisions. None of the other parameters were significantly different. mean±SD.

|  | **placebo** | **propranolol** | **amisulpride** | placebo vs propranolol |
| --- | --- | --- | --- | --- |
| $p_f$ | 21.01±4.47 | 16.52±6.27 | 18.86±5.88 | t(38)=2.60, p=.013 |
| $p_d$ | 13.90±5.89 | 9.14±4.23 | 11.77±6.38 | t(38)=2.94, p=.006 |
| $\tau_f$ | 2.85±2.23 | 3.30±2.47 | 3.26±2.58 | t(38)=-.60, p=.550 |
| $\tau_d$ | 5.10±2.74 | 4.73±2.35 | 5.52±2.68 | t(38)=.45, p=.653 |
| $\xi$ | 0.004±0.009 | 0.007±0.008 | 0.004±0.007 | t(38)=-1.21, p=.234 |

An important concomitant of an increasing urgency signal is a collapsing decision threshold. The threshold quantifies the amount of evidence needed at a particular stage in the information gathering process for committing to a decision. Its collapse indicates subjects adopt a more liberal decision criterion as more cards are turned over. Decision thresholds were significantly lower for the noradrenaline compared to the placebo group over most of the sampling process in both experimental conditions (Fig. 4, fixed: p=.036, decreasing p=.017).

**Figure 4.**
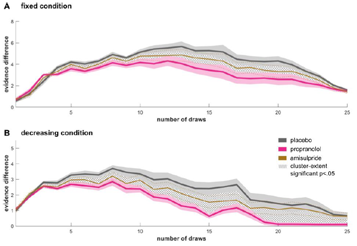
Lower decision threshold after noradrenaline blockade. A decision threshold (here captured as the evidence difference needed to make a decision at every stage) indicates how clear a choice must be for an agent to make a decision. This decision threshold shows a collapsing bound in both conditions, and is significantly lower in the noradrenaline (pink) compared to the placebo (gray) group in the fixed (A) and decreasing (B) condition. mean±1s.e.m.

## Discussion

An arbitration between gathering more information and time-efficiency is a non-trivial aspect of decision making, and it is thought to go awry in a range of psychiatric disorders such as schizophrenia or OCD (Fear and Healy, 1997; Moutoussis et al., 2011; Hauser et al., 2017c, 2017b). We show that inhibiting noradrenergic function by means of a noradrenergic beta-adrenoceptor blockade leads to decreased information gathering. Using computational modelling we show that noradrenaline blockade has a primary effect on a component of the decision process that reflects an ‘urgency to decide’. Urgency signals facilitate decision making, but have only been described in perceptual decision making (Cisek et al., 2009; Drugowitsch et al., 2012; Thura and Cisek, 2014; Murphy et al., 2016). Our results demonstrate that a similar urgency signal plays a role in sequential information gathering, where a decision urgency arises non-linearly.

Our computational modelling showed that noradrenaline directly modulates how quickly an urgency signal arises. Inhibiting noradrenaline function led to an earlier emergence of this urgency signal. Theories of noradrenaline functioning suggest that it influences cognition at different timescales, with tonic and phasic noradrenaline expressing distinct functional roles (Aston-Jones and Cohen, 2005; Yu and Dayan, 2005; Dayan and Yu, 2006). Because our manipulation is likely to influence both tonic and phasic noradrenaline, a specific contribution of one, or other, cannot be inferred from our data. We note a recent perceptual decision making study (Murphy et al., 2016) supports the idea that phasic noradrenaline might be critical for decision urgency. The authors showed that phasic pupil response, a potential indicator for phasic noradrenaline (Joshi et al., 2016), is linked to longer decision times possibly signalling a delayed emergence of an urgency signal. Whether such phasic noradrenaline promotes urgency directly, or whether it is mediated indirectly by signalling unexpected uncertainty (Dayan and Yu, 2006), which in turn delays an arising urgency, can be addressed in future studies.

Our findings suggest that manipulating noradrenaline could benefit patients whose information gathering is aberrant. In particular, we showed previously that compulsive subjects and patients with OCD gather information excessively and that this is due to a delayed emergence of decision urgency (Hauser et al., 2017c, 2017b). The antagonistic effect of propranolol on this urgency signal raises the theoretical possibility that this agent might alleviate an indecisiveness found in OCD patients. We are not aware of any study that has investigated the role of noradrenergic beta-receptor blockade in the treatment of OCD. Indirect evidence for the viability of this mode of intervention comes from observation with the therapeutic agent clomipramine. This is a first-line treatment for OCD (Koran et al., 2007) whose mode of action comprises complex noradrenergic effects including a down-regulation of beta-adrenoceptor density (Asakura et al., 1982), that might attenuate the impact of endogenous noradrenaline on beta-adrenoceptor related functions. Additionally, augmenting clomipramine with pindolol, a beta-blocker, is reported to have positive effects on OCD symptoms (Koran et al., 2007; Sassano-Higgins and Pato, 2015). These findings would support a more systematic examination of whether propranolol has beneficial effects for OCD patients in general, or more specifically patients where there is evidence for prepotent excessive information gathering behaviour.

Previous studies on the effects of dopaminergic drugs on sequential information gathering have produced mixed results. In schizophrenia, anti-psychotic medication does not directly alleviate jumping-to-conclusion behaviour (Menon et al., 2008; So et al., 2010, 2012), and even remitted patients express intermediate levels of information gathering (Moutoussis et al., 2011). In healthy subjects, only one study found a more cautious decision making after dopamine D2/3 agonist administration (Ersche et al., 2011), while several other studies did not find an effect of either dopamine enhancing or reducing medications (Andreou et al., 2013, 2015; Ermakova et al., 2014). Using amisulpride, we found no effect on information gathering. However, in almost all analysis, the effects of amisulpride were intermediate between placebo and propranolol, suggesting a similar, albeit weaker, effect than our noradrenaline manipulation. Whether this is due to a less direct influence of dopamine on information gathering, or whether it might be due to a lower effective dose of amisulpride (as compared to propranolol) remains unclear. However, the latter is less likely as there have been several studies showing significant effects of such a dose on cognition (e.g., Ramaekers et al., 1999; Kahnt and Tobler, 2017).

In conclusion, we show that information gathering is modulated by noradrenaline. Our computational modelling shows this is mediated via a modulation of a subjective, non-linear, urgency-to-respond signal. An involvement of noradrenaline in this aspect of decision making opens potential avenues for therapy in psychiatric conditions where there is aberrant information gathering.

## Acknowledgements

The Wellcome Trust’s Cambridge-UCL Mental Health and Neurosciences Network grant (095844/Z/11/Z) supported RJD, TUH and MM. TUH is supported by the Jacobs Foundation. RJD holds a Wellcome Trust Senior Investigator Award (098362/Z/12/Z). MM was also supported by the Biomedical Research Council. PD was supported by the Gatsby Charitable Foundation. The Max Planck UCL Centre is a joint initiative supported by UCL and the Max Planck Society. The Wellcome Trust Centre for Neuroimaging is supported by core funding from the Wellcome Trust (091593/Z/10/Z). PD is currently on sabbatical, working at Uber Technologies, but the company had not influence on this work. The other authors declare no competing financial interests.

